# Monoclonal Antibodies from COVID-19 Convalescent Patients Target Cryptic Epitopes for Universal SARS-CoV-2 Neutralization

**DOI:** 10.1101/2025.08.17.670707

**Authors:** Aakanksha Harit, Michael Mor, Ron Yefet, Lee S. Izhaki-Tavor, Meital Gal-Tanamy, Natalia T Freund, Moshe Dessau

## Abstract

The COVID-19 pandemic, which has resulted in over seven million global fatalities, poses a substantial threat to public health and precipitated a global economic crisis. Emerging variants of concern (VOCs) with enhanced transmissibility and improved immune evasion may compromise the efficacy of current antiviral and immunotherapies, necessitating comprehensive investigations into the immune response to SARS-CoV-2. The conformational dynamics of the receptor binding domain (RBD) in SARS-CoV-2 spike and the presentation of neutralizing antibody epitopes influence viral transmission and infection rates. In this study, we have identified highly conserved non-RBM epitopes for two potent monoclonal antibodies (mAbs), TAU-1109 and TAU-2310, isolated from convalescent human patients, which contribute to the broad neutralizing activity of these mAbs against all the circulating VOCs, including the recently emerged Omicron subvariants. We employed high-resolution structural data in conjunction with systematic biochemical investigation to elucidate the neutralization mechanism of TAU-1109 and TAU-2310. The mechanism involves antibody-mediated destabilization of the spike trimer, resulting in the premature shedding of the S1 subunit and rendering the spike incapable of mediating host cell entry. The identification of conserved cryptic epitopes in our study advances the mechanistic understanding of immune response against SARS-CoV-2, providing novel avenues for the development of universal therapeutic antibodies and vaccines to combat COVID-19.

## Introduction

The COVID-19 pandemic, caused by the newly emerged severe acute respiratory syndrome coronavirus 2 (SARS-CoV-2), has led to over 778 million confirmed human infections globally as of July 2025 (https://covid19.who.int/). The continued global spread of the virus has been accompanied by the emergence of new variants of concern (VOCs) with enhanced transmissibility and immune evasion capabilities. These new VOCs have completely displaced the original Wuhan-Hu-1 strain in subsequent waves of the pandemic. Their emergence poses a significant threat to the effectiveness of current antiviral and immunotherapeutic regimens, underscoring the urgent need to design and develop new vaccines through a detailed investigation of the immune response against SARS-CoV-2.

SARS-CoV-2 is an enveloped, positive-sense, single-stranded RNA virus classified under the *Betacoronavirus* genus, which also includes SARS-CoV, MERS-CoV, and several other coronaviruses known to infect humans (e.g., HCoV-OC43, HCoV-HKU1) and various other animal species ^1^. The SARS-CoV-2 encoded spike (S) glycoprotein is responsible for viral recognition and attachment to the host cell receptor, the human angiotensin-converting enzyme 2 (ACE2) ^2^. The spike protein assembles as homotrimers on the viral membrane, where each monomer comprises two subunits, S1 and S2, with distinct functions. Membrane fusion is a critical step for establishing infection by enveloped viruses. It requires a carefully orchestrated structural rearrangement of the viral envelope glycoproteins, refolding the metastable pre-fusion state into a stable post-fusion conformation, thereby overcoming the high kinetic energy barriers involved in fusing the viral and host cell membranes and enabling delivery of the viral genome into the host cytoplasm ^3^. In the pre-fusion state, the inherently dynamic S1 subunit, containing the N-terminal domain (NTD) and receptor- binding domain (RBD), mediates the binding to ACE2, SARS-CoV-2’s cellular receptor, while the S2 subunit acts as a stalk anchoring the S protein to the virion membrane. During host cell entry, RBD-ACE2 binding and proteolytic activation of spike by the human protease TMPRSS2 trigger the dissociation (shedding) of the S1 subunit, and a conformational change in the S2 subunit exposes the fusion peptide (FP), which mediates the fusion of the viral and cellular membranes ^4,5^. The RBD transitions between two main conformations, representing a receptor-inaccessible (‘closed’ or ‘down’) and receptor-accessible (‘open’ or ‘up’) state ^6^. Hence, the conformational dynamics of the RBD dictate epitope accessibility to neutralizing antibodies and play a crucial role in SARS-CoV-2 evolution by influencing viral transmission and immune evasion ^7^.

The severity of SARS-CoV-2 infection can clinically range from asymptomatic carriers to severe disease, resulting in high morbidity and mortality in affected individuals. The interaction between SARS-CoV-2, neutralizing antibodies, and immune cells contributes to both pathogenesis and protective immunity against the disease ^8^. Severe disease can be prevented either by the development of an effective humoral immune response that limits viral dissemination or by T-cell-mediated immunity that restricts viral replication and disease progression ^9^. Therefore, studying neutralizing antibodies that target the spike glycoprotein has been crucial for advancing our understanding of the immune response against SARS-CoV-2. A wide range of neutralizing antibodies (nAbs) have been identified, each recognizing distinct epitopes on the S1 and S2 subunits of the spike protein ^10–16^. Many of the reported neutralizing antibodies target the RBD within the S1 subunit ^17^. Until recently, RBD-targeting antibodies were classified into four primary classes based on their epitopes and neutralization mechanisms ^18^. Classes 1 and 2 specifically recognize epitopes that overlap with the receptor-binding motif (RBM) ^10,19^. However, many of these nAbs do not retain activity against later VOCs as they target mutation-sensitive epitopes ^20^. Class 3 antibodies bind outside the ACE2 binding site, providing cross-variant protection ^12^, while class 4 antibodies target more conserved residues, sterically hindering ACE2 binding ^11^. Notably, newly emerging Omicron subvariants significantly evade all four classes of RBD-targeting antibodies, resulting in immune evasion ^21^. Recently, two novel classes of RBD-targeting antibodies (classes 5 and 6) were described, each recognizing highly conserved and cryptic epitopes outside the RBM and demonstrating potential broad neutralization capabilities ^22,23^.

Here, we investigate the structural and neutralization mechanism of two anti-SARS-CoV-2 antibodies (TAU-1109 and TAU-2310) isolated from severe convalescent donors among the first ten COVID-19 cases documented in Israel, who were infected with the wild-type Wuhan-Hu-1 strain in March 2020 ^24^. Both antibodies demonstrated neutralization activity against lentivirus-based pseudo-virus particles presenting SARS-CoV-2 spike on their membrane; however, only TAU-1109 was able to neutralize the authentic wild-type SARS-CoV-2 virus when introduced to Vero E6 cells^25^. Interestingly, both antibodies retain their efficacy against other SARS-CoV-2 VOCs, including Alpha, Beta, Gamma, Delta, and Omicron, indicating that most mAbs targeting non-ACE2 binding sites are less sensitive to viral mutations. Comparison of B-cell receptor (BCR) signatures from severe patient donors with BCR repertoires from unexposed naïve and mature B cells shows the presence of precursor antibodies for these nAbs, suggesting that such antibodies can be readily produced by the majority of the unexposed population upon antigenic stimulation ^24^

Here, we employed X-ray crystallography to delineate the RBD epitopes recognized by the monoclonal antibodies TAU-1109 and TAU-2310. The latter targets a highly conserved epitope characteristic of class 6 antibodies. In contrast, structural analysis revealed that TAU-1109 engages an atypical epitope overlapping features of both class 5 and class 6 antibodies, with an unusually extensive footprint. Notably, TAU- 1109 effectively competes with antibodies targeting this region, including TAU-2310. Collectively, our findings provide structural and functional insights into the modes of action of broadly cross-neutralizing anti-SARS-CoV-2 antibodies and offer a valuable framework for the rational design of next-generation antibody-based therapeutics and vaccines.

## Methods

### Cell lines

*Drosophila* S2 cell lines were cultured in ESF921 protein-free medium (Expression Systems), suspended at 27°C with shaking at 110 rpm. Expi293F cells (Thermo- Fisher) grew in Expi293^TM^ Expression Medium (ThermoFisher) at 37°C and 8% CO2, shaking at 130 rpm. The HEK 293T and HEK 293T-hACE2 cells were grown in DMEM (Sartorius) at 37°C and 5% CO2.

### Sub-cloning, protein expression, and purification of soluble wild-type SARS-CoV-2 RBD

#### Insect Cell Expression

The RBD of wild-type SARS-CoV-2 spike (residues 319-541) was subcloned into pMT-BiP-His-C (Invitrogen) in frame with a C-terminal HRV 3C protease site and cleavable 2×StrepTag (pMT-wtRBD). *Drosophila* S2 cells were used to generate stable transfectants of the pMT-wtRBD construct. The cells were co-transfected with pMT-wtRBD plasmid and pCoPuro (puromycin resistance) plasmid in a ratio of 20:1 using ESCORT IV transfection reagent (Sigma Aldrich). The cells were then selected with 7 μg/ml puromycin (InvivoGen) for 2-3 weeks until stable cells regain exponential growth. Stably transfected cells were frozen at 1×10^7^ cells/ml and stored in LN2 until further use. The expression of wild-type RBD was induced using 600 μM CuSO4 solution at a cell density of 1×10^7^ cells/ml. After 6 days, cells were discarded, and the S2 media supernatant was concentrated to 150 ml while exchanging the buffer to 100 mM Tris-HCl, 150 mM NaCl, 1 mM EDTA pH 8.0 (binding buffer) using a tangential flow filtration (TFF) system (PALL). The sample was then centrifuged at 30,000×g/30 min/4°C (Avanti JE Centrifuge) and loaded onto strep- tactin affinity column (GE Healthcare). The protein was eluted with 2.5 mM desthiobiotin (Sigma Aldrich). The strep-tag was removed by an overnight digestion with HRV 3C protease (1:100 molar ratio) at 4°C and further purified on a size-exclusion chromatography column (Superdex 200, GE Healthcare) pre-equilibrated with 20 mM Tris pH 8.0 and 150 mM NaCl.

#### Mammalian Cell Expression

The wtRBD (residues 311-519) was subcloned into pHL-sec vector containing a native secretory signal (MGILPSPGMPALLSLVSLLSVLLMGCVA) and CMV enhancer between AgeI and KpnI restriction sites, in frame with a C-terminal cleavable 6×Histag. The Expi293F cells (ThermoFisher) were transiently transfected with the wtRBD plasmid using ExpiFectamine 293 Transfection Kit (ThermoFisher) according to the manufacturer’s instructions. Five days post-transfection, the media (supernatant) were collected and purified using Ni Sepharose 6 resin (Cytiva).

#### Cloning, protein expression, and purification of IgGs and Fab fragments

The heavy and light chain residues of the TAU-1109 and TAU-2310 mAbs were subcloned in the pCMV vector backbone downstream of a murine Ig signal peptide. The Expi293F (ThermoFisher) cells were co-transfected with the heavy chain and light chain of TAU- 1109 and TAU-2310 in the ratio of 1:1 using ExpiFectamine 293 Transfection Kit (ThermoFisher) according to the manufacturer’s instructions. Five days post-transfection, the media (supernatant) was collected, and the media condition was adjusted to the binding buffer (20 mM sodium phosphate pH 7). The filtered supernatant was purified using HiTrap protein G HP antibody purification column (GE Healthcare). The protein was eluted with 0.1 mM glycine-HCl pH 2.7, followed by neutralization with 1M Tris pH 9. The elution fractions were analyzed using SDS-PAGE, pooled, and subjected to a size-exclusion chromatography column (Superdex 200, GE Healthcare) pre-equilibrated with 20 mM HEPES pH 8.0 and 150 mM NaCl.

For the TAU-1109 and TAU-2310 Fab fragments, the variable region of the heavy chains was subcloned into the phCMV1-CH1 backbone using Gibson Assembly (NEB). The Expi293F (ThermoFisher) cells were co-transfected with the Gibson-cloned Fab heavy chain and light chain of TAU-1109 and TAU-2310 in the ratio of 1:1 using ExpiFectamine 293 Transfection Kit (ThermoFisher). The protein was purified similarly to that described for the purification of TAU-1109 and TAU-2310 mAbs.

A broad sarbecovirus neutralizing mAb (5817) ^26^ was used as a structural and functional comparison. Briefly, the IgH/IgL sequences of 5817 were obtained from the same source ^26^, and the corresponding amino acid sequences were codon-optimized, synthesized by an external vendor, and cloned into human IgG1 and IgL immunoglobulin expression vectors. The cloned plasmids were transiently co-transfected at a 1:3 ratio (respectively) into Expi293F cells, following the manufacturer’s protocol. Seven days post-transfection, the supernatant was incubated with protein A beads (Cytiva) for 2 hours at room temperature (RT). Beads were then transferred to chromatography columns, washed, and eluted using a low-pH buffer, followed by overnight dialysis against PBS. Proper production of the 5817 mAb was verified by testing its binding to the WT RBD in ELISA before its subsequent use.

#### Cloning, protein expression, and purification of RBD mutants based on the TAU- 1109 epitope

Site-directed mutagenesis was performed using our previously described wild-type plasmid ^24^ as a template to generate RBD variants containing amino acid substitutions. Overlapping primer pairs carrying one or two base changes, flanked by ∼20 nucleotides on either side, were designed and synthesized by Syntezza (Israel) (Table S4). PCR amplification was carried out with KAPA HiFi HotStart ReadyMix (Roche), using 10 µL of the mix, 0.5 µM of each primer, and 1 ng of plasmid template in a final volume of 20 µL adjusted with DNase/RNase-free water (Bio-Lab). The thermal cycling program consisted of an initial denaturation at 95 °C for 3 min, followed by 16 cycles of 98 °C for 20 s and 72 °C for 90 s. Multiple substitutions were introduced by repeating the procedure with appropriate mutant templates and primers. The resulting constructs were transiently transfected into Expi293F cells (Thermo Fisher) using the ExpiFectamine 293 Transfection Kit according to the manufacturer’s protocol. Seven days after transfection, supernatants were harvested, clarified by 0.22 µm filtration, and incubated with Ni²⁺-NTA agarose beads (GE Life Sciences) for 2 h at room temperature. Bound proteins were eluted with 200 mM imidazole, buffer-exchanged into 1× PBS, aliquoted, and stored at −80 °C.

### Cloning, protein expression, and purification of RBD mutants based on the TAU- 2310 epitope

To generate RBD mutants corresponding to the TAU-2310 epitope, site-directed PCR mutagenesis was performed using the pHL-sec_wtRBD_6×His construct as a template. The primer sequences are given in Table S4. The constructs were verified using Sanger sequencing.

### Purification, crystallization, and structure determination of RBD-Fab immune complexes

The purified Fab fragments of TAU-1109 and TAU-2310 were mixed with wtRBD in a 1:1 molar ratio, respectively, and incubated for 1 hour on ice. The resultant complexes were purified using size-exclusion chromatography (Superdex 200, GE Healthcare) and eluted with 20 mM HEPES pH 8.0, and 150 mM NaCl. The relevant peak fractions of the respective complexes were collected for crystallization.

The crystallization trials were performed at 16°C using the hanging-drop vapor-diffusion method with different commercial crystal screens in a 2:1 protein-to-reservoir ratio using the Mosquito Nanodrop Crystallization Robot (TTP Biotechnology).

For the RBD-Fab^TAU-1109^ complex, initial tiny rod-shaped crystals appeared after 16-18 weeks in wells containing 0.2 M zinc acetate dihydrate, 0.1 M sodium cacodylate trihydrate pH 6.5, and 18% (w/v) PEG 8000. The crystallization condition was further optimized by varying pH and precipitant concentration to get diffraction-quality crystals. In the optimization screen, multiple rod-shaped crystals appeared in 5-6 weeks in 0.2 M zinc acetate dihydrate, 0.1 M sodium cacodylate trihydrate pH 5.5, and 15% PEG 8000. The crystal growth took another 2-3 weeks before the final data collection. The crystals were cryoprotected in the reservoir solution supplemented with 15% glycerol.

For the RBD-Fab^TAU-2310^ complex, thin plate-like crystals appeared after 2 days in wells containing 0.2 M lithium citrate tribasic tetrahydrate and 20% (w/v) PEG 3350. The crystals were cryoprotected in the reservoir solution supplemented with 24% glycerol. The X-ray diffraction data were collected at the European Synchrotron Radiation Facility (ESRF) at ID30B and ID23-2 beamlines for RBD-Fab^TAU-1109^ and RBD-Fab^TAU-2310^ complexes, respectively. The crystals of RBD-Fab^TAU-1109^ belong to an orthorhombic lattice and are indexed to the *I 212121* space group. The structure was determined by molecular replacement with PDB 7E3O as a search model using PHASER, where the RBD and the Fab were used as two separate search models. The model obtained in the molecular replacement was rebuilt according to the TAU- 1109 sequence using COOT, and the structure was refined with PHENIX to 2.56 Å resolution with excellent statistics. The crystals of RBD-Fab^TAU-2310^ belong to a monoclinic lattice and are indexed to the *C2* space group. The structure was determined by molecular replacement with AlphaFold 3 prediction as a search model using PHASER, where the RBD and the Fab were used as two separate search models. The structure was refined with PHENIX to 2.12 Å resolution with excellent statistics. The data collection and refinement statistics are listed in Table 1. The atomic coordinates and structure factors for the RBD-Fab^TAU-1109^ and RBD-Fab^TAU-2310^ immune complexes were deposited to the RCSB PDB under accession codes 9SAT and 9SBB, respectively. All molecular graphics images are produced using PyMol (PyMOL Molecular Graphics System, Version 2.1, Schrödinger, LLC).

**Table 1.** Data collection and refinement statistics.

|  | wtRBD-Fab<sup>TAU-1109</sup> complex | wtRBD-Fab<sup>TAU-2310</sup> complex | 646 |
| --- | --- | --- | --- |
| **Data Collection statistics** |  |  |  |
| PDB ID | 9SAT | 9SBB | 647 |
| Wavelength | 0.8731 | 0.8731 |  |
| Resolution range | 93.7 - 2.56 (2.65 - 2.56) | 60.42 - 2.119 (2.16 - 2.12) | 648 |
| Space group | *I* 2<sub>1</sub> 2<sub>1</sub> 2<sub>1</sub> | *C* 1 2 1 |  |
| Unit cell. a, b, c | 83.3 103.0 225.5 | 111.8 88 95.8 |  |
| α, β, γ | 90 90 90 | 90 108.25 90 | 649 |
| Total reflections | 55264 (5129) | 169536 (9313) |  |
| Unique reflections | 28734 (2658) | 49442 (2730) | 650 |
| Multiplicity | 1.9 (1.9) | 3.4 (3.4) |  |
| Completeness (%) | 90.72 (85.56) | 98.60 (91.92) |  |
| Mean I/sigma(I) | 9.39 (3.65) | 7.52 (0.84) | 651 |
| Wilson B-factor | 21.78 | 35.51 |  |
| †R-merge | 0.067 (0.232) | 0.13 (1.625) | 652 |
| R-meas | 0.095 (0.329) | 0.1544 (1.926) |  |
| R-pim | 0.067 (0.232) | 0.08232 (1.021) |  |
| CC1/2 | 0.99 (0.84) | 0.993 (0.372) | 653 |
| CC* | 0.99 (0.96) | 0.998 (0.737) |  |
| **Refinement statistics** |  |  | 654 |
| Reflections used in refinement | 28727 (2654) | 49432 (2732) |  |
| Reflections used for R-free | 1416 (142) | 2449 (120) | 655 |
| R-work | 0.2309 (0.2881) | 0.1968 (0.3536) |  |
| R-free | 0.2753 (0.3597) | 0.2302 (0.3997) |  |
| Number of non-hydrogen atoms | 5098 | 4998 | 656 |
| macromolecules | 4843 | 4741 |  |
| ligands | 3 | 40 | 657 |
| solvent | 252 | 217 |  |
| Protein residues | 627 | 624 |  |
| RMS(bonds) | 0.01 | 0.01 | 658 |
| RMS(angles) | 1.46 | 1.47 |  |
| Ramachandran favored (%) | 95.8 | 97.08 | 659 |
| Ramachandran allowed (%) | 4.2 | 2.92 |  |
| Ramachandran outliers (%) | 0 | 0 |  |
| Rotamer outliers (%) | 2.22 | 0.76 | 660 |
| Clashscore | 8.72 | 5.65 |  |
| Average B-factor | 33.05 | 48.25 | 661 |
| macromolecules | 33.37 | 48.10 |  |
| ligands | 36.45 | 81.75 |  |
| solvent | 26.97 | 45.48 | 662 |

### Analysis of epitope conservation and antigen-antibody contacts

The conservation of the epitopes was determined using the ConSurf web server ^27^. The figure indicating the scale of sequence conservation was generated using PyMOL (PyMOL Molecular Graphics System, Version 2.1, Schrödinger, LLC). The structural analysis of the RBD and Fabs was performed using CCP4i. The Van der Waals contacts having an interatomic distance of ≤ 4Å were defined. The putative hydrogen bonds were defined by PISA.

### Lentivirus-based pseudo-particle preparation and neutralization assays

To generate Lentivirus-based vectors pseudotyped with spike (S) glycoproteins of different VOCs, Expi293F (ThermoFisher) cells were co-transfected with pCMV delta R8.2, pLenti GFP (GeneCopoeia), and pcDNA3.1 SΔC19 (or spike glycoproteins of respective VOCs) at a ratio of 1:2:1, respectively, according to the manufacturer’s instructions. The supernatant was harvested 72 hours post-transfection, centrifuged at 1500×g for 10 minutes, filtered, aliquoted, and stored at -80°C until further use.

For the neutralization assays, HEK 293T cells stably expressing hACE2 were seeded into poly-D-lysine-coated 96-well plates (Greiner) at an initial density of 0.4x10^5^ cells per well. The following day, the lentiviral particles were concentrated to 10% of their original volume using an Amicon^â^ Ultra centrifugal filter of 100kDa cutoff (Merck Millipore). The concentrated pseudo-particles (at 0.1 MOI) were then incubated with serial dilutions of TAU-1109 and TAU-2310 for 1 hour at 37°C and added to the pre-seeded 96-well plates. After 24 hours, the plates were imaged with a 10x objective using the IncuCyte SX5 system (Sartorius) to calculate GFP-positive cells from four representative images per well (data for each antibody was collected in triplicate). The number of GFP-positive cells was normalized to calculate relative infection (%) and plotted against the antibody dilutions (μg/ml) using GraphPad Prism (v10.4.1). The IC50 values (μg/ml) were calculated using GraphPad Prism (v10.4.1).

The lentiviral particles expressing spike glycoproteins of Wuhan-Hu-1 and Omicron subvariants were produced and tested similarly. The plasmids pcDNA3.3_SARS2_BQ.1.1 (Addgene plasmid # 194493) and pcDNA3.3_SARS2_XBB.1.5 (Addgene plasmid # 196585) were kind gifts from David Nemazee.

### Spike S1 shedding assay

HEK 293T cells were transfected with pcDNA 3.1 SΔC19 plasmid using PEI (Sigma Aldrich) and incubated for 48 hours at 37°C and 5% CO2. The cells were resuspended in FACS buffer (ice-cold PBS and 2% Fetal Bovine Serum). The TAU-1109/TAU-2310 (100 nM) was serially incubated with cells for different time points (120, 60, and 5 minutes) on ice. The plates were thoroughly washed twice with FACS buffer and stained with donkey anti-human IgG H&L (DyLight^â^ 488) (Abcam) (1:100 dilution) on ice for 30 minutes (in the dark). After two washes with FACS buffer, the cell samples were resuspended and analyzed using Gallios Flow Cytometer (Beckman Coulter) and FlowJo software. The mean fluorescence intensity (MFI) for each sample was determined at each time point, and each sample was normalized to the MFI at the 5 min time point (MFI/MFI 5 min×100).

For western blot, the spike-transfected HEK 293T cells were resuspended in Dulbecco’s Phosphate Buffer Saline (Sartorius) and were serially incubated with 100 nM of TAU-1109 and TAU-2310 for different time points (120, 60, and 5 minutes) on ice. After incubation, the cell lysates and supernatants were subjected to Western blotting using SARS-CoV-2 spike protein S1 recombinant Rabbit monoclonal antibody (HL6) (Invitrogen) (1:5000), followed by Goat anti-Rabbit HRP conjugated secondary antibody (1:10,000). The bands were visualized with Amersham ECL Western Blotting Detection Reagent (Cytiva) using Platinum Q9 (Uvitec) gel documentation system.

### BioLayer Interferometry (BLI)

The BLI experiments were performed on an Octet Red96e instrument (ForteBio) at 30°C with shaking at 1,000 RPM. The TAU-1109 and TAU-2310 mAbs were diluted in Kinetics Buffer (1x PBS pH 7.4 and 0.02% Tween 20) to a concentration of 10 μg/ml and immobilized onto Anti-Human IgG Fc Capture (AHC) Biosensors (ForteBio) for 300s. The sensors were then immersed in the kinetics buffer for 60s to record a baseline. Purified SARS-CoV-2 RBDs of wild-type and different Omicron subvariants were serially diluted (200, 100, 50, 25, 12.5, 6.25, 3.125 nM) in kinetics buffer, and an association was recorded for 600s. The dissociation was then measured by immersing the sensors in kinetics buffer for 600s to perform the kinetic analysis using the global 1:1 binding curve fitting model in ForteBio data analysis HT software (Version 11.1.0.25). *K*D, *k*on, *k*off values were determined by averaging all binding curves that matched the theoretical fit with an R^2^ value of ≥ 0.95. The kinetic graphs were plotted using GraphPad Prism (v10.4.1).

For affinity analysis of RBD mutants, 200 nM of wild-type RBD and respective mutants were associated with TAU-1109/TAU-2310 (10 μg/ml) loaded Anti-Human IgG Fc Capture (AHC) Biosensors for 600s, followed by a 600s dissociation in kinetic buffer. The kinetic analysis was performed using a local (individual) 1:1 binding curve fitting model in ForteBio data analysis HT software (Version 11.1.0.25). The kinetic graphs were plotted using GraphPad Prism (v10.4.1).

For the competition assay, biotinylated wild-type SARS-CoV-2 RBD (10 μg/ml) was immobilized onto High Precision Streptavidin (SAX) Biosensors (Sartorius) for 300 seconds. The sensors were then immersed in 1x PBS for 60 seconds to record a baseline. TAU-1109 (200 nM) and TAU-2310 (200 nM) antibodies were sequentially associated for 600 seconds each after a 60 seconds baseline step. The curves were plotted using GraphPad Prism (v10.4.1). Similar experiments were performed for the competition analysis of TAU-1109 and TAU-2310 with Ab 5817, respectively.

## Results

### Measuring the binding affinities and neutralization breadth of TAU-1109 and TAU-2310 antibodies against SARS-CoV-2 Variants of Concern (VOCs)

The rapid emergence of diverse SARS-CoV-2 variants has compromised the efficacy of many therapeutic antibodies ^28–30^, highlighting the need to identify broadly neutralizing antibodies for developing pan-coronavirus vaccines. TAU-1109 and TAU- 2310, derived from two distinct B cell clones (VH1-18 and VH3-23, respectively), were isolated from the PBMCs of two different severe convalescent donors amongst the first ten documented COVID-19 cases in Israel, who were initially infected with the wild-type Wuhan-Hu-1 strain ^24^. Previous studies demonstrated that these mAbs bind to the SARS-CoV-2 RBD and stabilized spike trimers ^25^. To further characterize their binding properties, we performed quantitative kinetic analyses to determine the association and dissociation constants of TAU-1109 and TAU-2310 against various SARS-CoV-2 VOCs. Both antibodies exhibited strong binding affinities and broad reactivity toward the RBDs of the wild-type Wuhan-Hu-1 strain, Omicron, and its sub- lineages, with KD values ranging from sub-nanomolar to nanomolar levels, confirming their classification as high-affinity mAbs (Fig.1A, Table S1). TAU-1109 demonstrated better stability (low *k*dis) and higher *k*a across different strains. In contrast, TAU-2310 showed faster association rates (*k*a) for early Omicron variants (B.1.1.529, BA.2, and BA.2.12.1), and slower dissociation against newer variants such as XBB.1.5.

**Fig 1.**
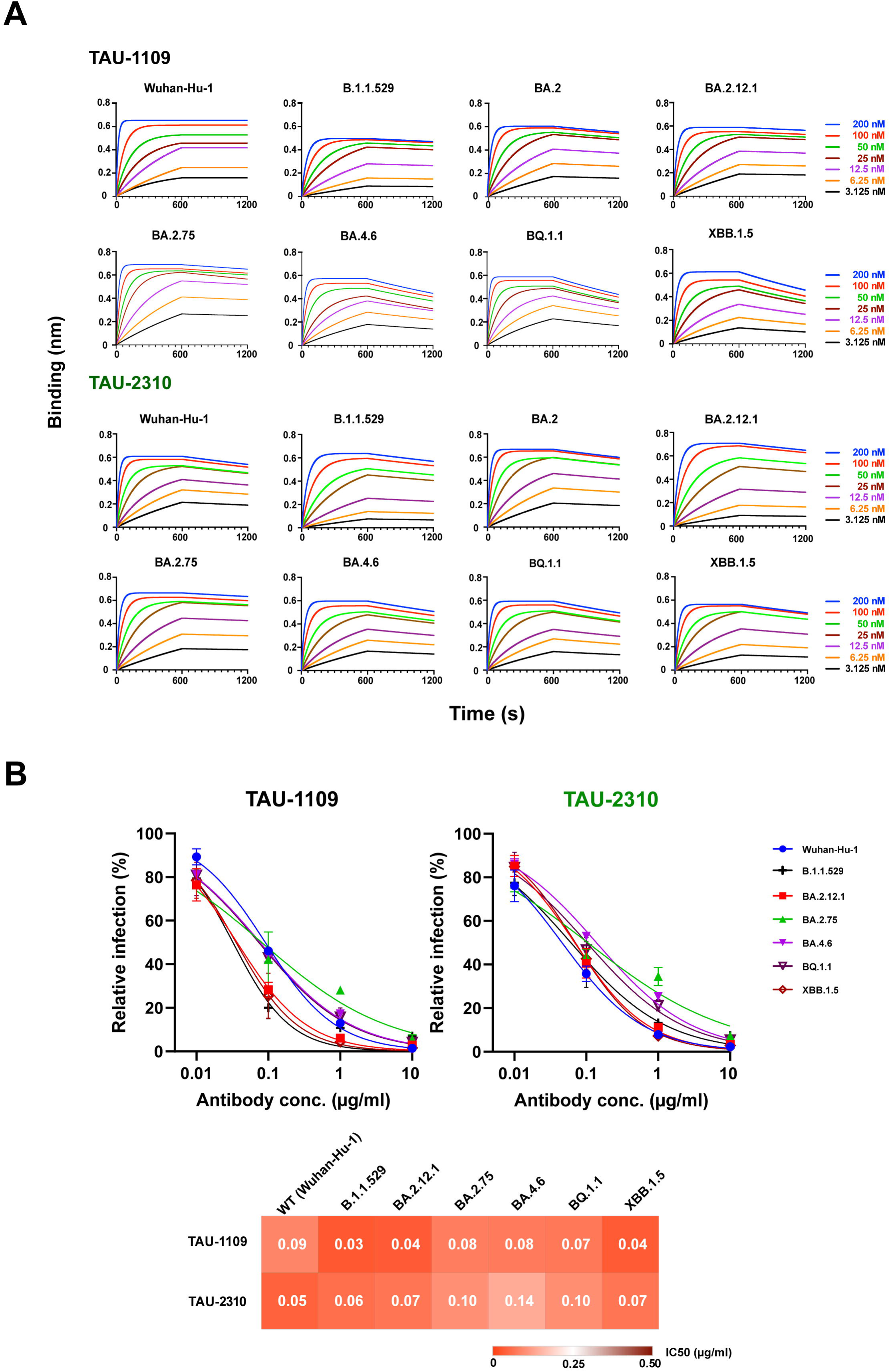
**Characterization of binding affinity and neutralization breadth of TAU-1109 and TAU-2310 mAb** A) BioLayer Interferometry (BLI) was used to conduct kinetic analyses to determine the binding affinities of TAU-1109 and TAU-2310 against RBDs of different VOCs of SARS-CoV-2 on the Octet Red96e instrument (ForteBio) using Anti-Human IgG Fc Capture (AHC) Biosensors (ForteBio) to capture antibodies, followed by association with respective RBDs. The kinetic graphs were plotted using GraphPad Prism (v10.4.1). B) Neutralization efficiencies of TAU-1109 and TAU-2310 against various VOCs were tested by the Lentivirus-based pseudovirus system. The mixture of lentiviral particles and serially diluted mAbs was added to HEK293T-hACE2 cells for 24 hours. The GFP-positive cells were imaged with a 10x objective using the IncuCyte SX5 system (Sartorius). The IC50 values (μg/ml) were calculated using GraphPad Prism (v10.4.1).

It was previously reported that TAU-1109 and TAU-2310 can neutralize the early SARS-CoV-2 VOCs, including the Omicron (B.1.1.529) variant ^25^. To determine the neutralization efficiency of these antibodies against newly emerged Omicron subvariants, we performed lentivirus-based neutralization assays (Fig. 1B). Both TAU- 1109 and TAU-2310 efficiently neutralized the Wuhan-Hu-1 strain and newly emerged Omicron subvariants, demonstrating potent cross-neutralization with comparable IC50 values (Fig. 1B). The low IC50 values observed for both mAbs indicate high potency against all circulating SARS-CoV-2 VOCs, including the immune-evasive Omicron sublineages.

### The cross-neutralizing properties of TAU-1109 and TAU-2310 are attributed to their recognition of highly conserved epitopes on RBD

Since the onset of the COVID-19 pandemic, the emergence of SARS-CoV-2 variants has challenged the efficacy of existing neutralizing antibodies (nAbs), but TAU-1109 and TAU-2310, isolated from convalescent patients, have shown cross-neutralizing potential against Omicron subvariants, likely due to their targeting of conserved RBD residues without competing with ACE2 binding ^24,25^. To elucidate the structural basis of TAU-1109 and TAU-2310 recognition of the RBD, we determined the crystal structures of the RBD-Fab^TAU-1109^ and RBD-Fab^TAU-2310^ immune complexes.

The RBD-Fab^TAU-1109^ complex was crystallized in 0.2 M zinc acetate dihydrate, 0.1 M sodium cacodylate trihydrate pH 6.5, and 18% (w/v) PEG 8000. Crystals were observed 5-6 weeks post optimizations and belonged to the *I 212121* orthorhombic space group. The structure was determined and refined at 2.56 Å (Table 1). The asymmetric unit of the crystal encompasses one RBD-Fab^TAU-1109^ complex (Fig. 2A). The RBD-Fab^TAU-1109^ interface, with buried surface area of about 1902 Å^2^, is composed of the residues Y351, A352, W353, N354, R355, K356, R357, S359, N360, T393, N394, Y396, P463, F464, E465, R466, I468, S469, T470, E471, F486, F490, L492, E516, L518, and A520 in the RBD (Fig. 2B). The paratope of TAU-1109 mAb mainly composed of complementarity-determining region (CDR) loops, with all six CDRs participating in the extended surface of the interaction: CDRH1 (residues 26-33), CDRH2 (residues 51-58), CDRH3 (residues 100-111), CDRL1 (residues 27-32), CDRL2 (residues 49-51), and CDRL3 (residues 89-98) (Fig. 2B). The immune complex is stabilized by 14 direct hydrogen bonds, contributed by RBD residues R355, R357, S359, P463, F464, R466, T470, E471, and F486, as well as Van-der-Waals contacts, as depicted in Fig. 3A-B, and summarized in Table 2.

**Fig 2.**
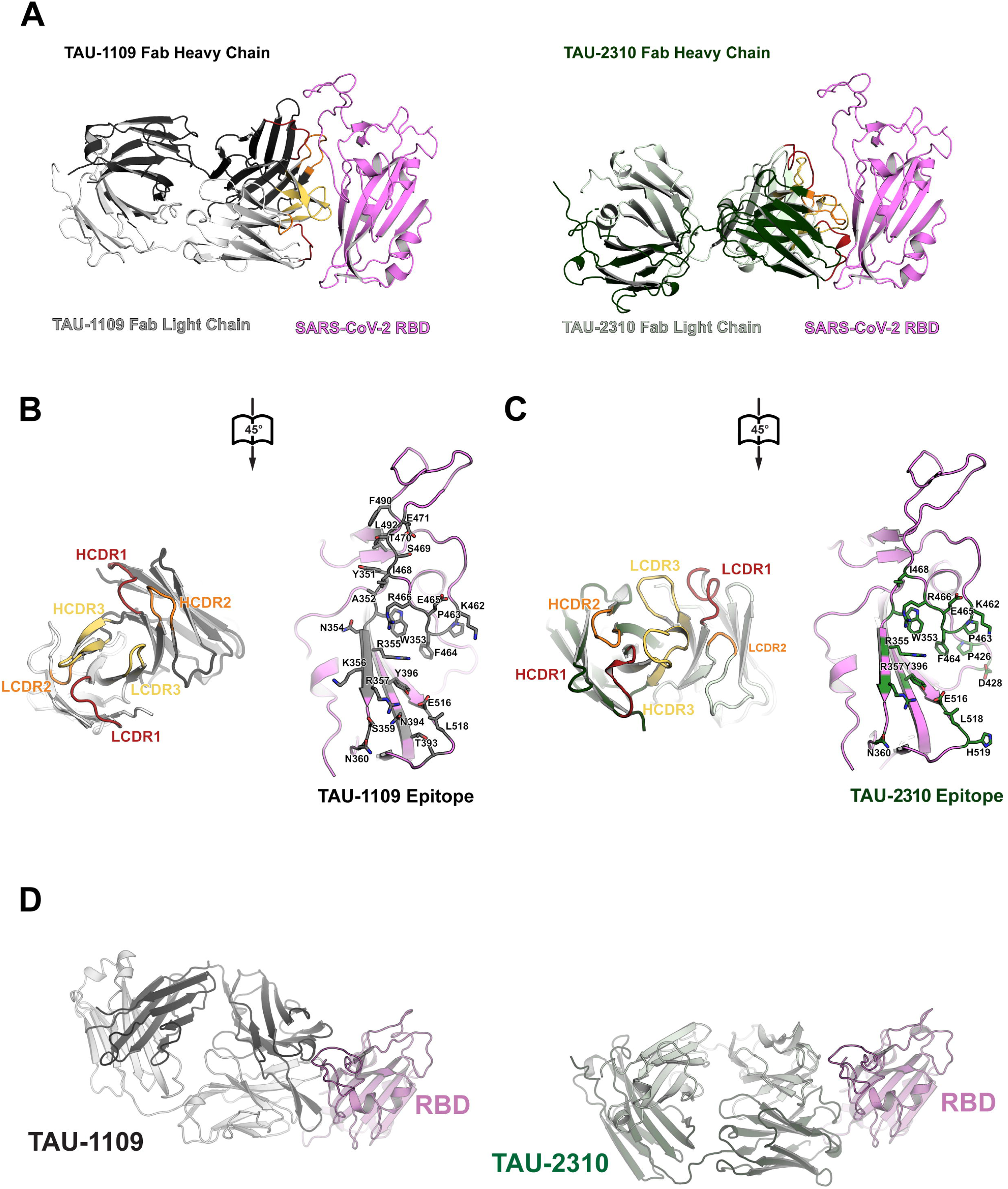
**TAU-1109 and TAU-2310 recognize unique epitopes on the SARS-CoV-2 receptor-binding domain** A) Cartoon representation of the SARS-CoV-2 RBD-Fab^TAU-1109^ (left) and RBD-Fab^TAU-2310^ (right) immune complexes. The color scheme displays WTRBD, TAU-1109 Fab HC, TAU-1109 Fab LC, TAU-2310 Fab HC, and LC in violet, gray 30, gray 80, forest green, and sick green, respectively. B) Open-book representation of the TAU-1109 epitope and the paratope. The interface residues on the SARS-CoV-2 RBD (in violet) are shown in gray 50 and are labeled accordingly. The complementarity-determining regions (CDRs) I, II, and III are shown in red, orange, and yellow, respectively. C) Open-book representation of the TAU-2310 epitope and the paratope. The interface residues on the SARS-CoV-2 RBD (in violet) are depicted in forest green and are labeled accordingly. The complementarity-determining regions (CDRs) I, II, and III are represented in red, orange, and yellow, respectively. D) Binding comparison of TAU-1109 and TAU-2310 to the SARS-CoV-2 RBD. TAU- 1109 targets a wider area on the RBD. Most interactions in the RBD-Fab^TAU-2310^ immune complex are mediated by the 2310Fab heavy chain.

**Fig 3.**
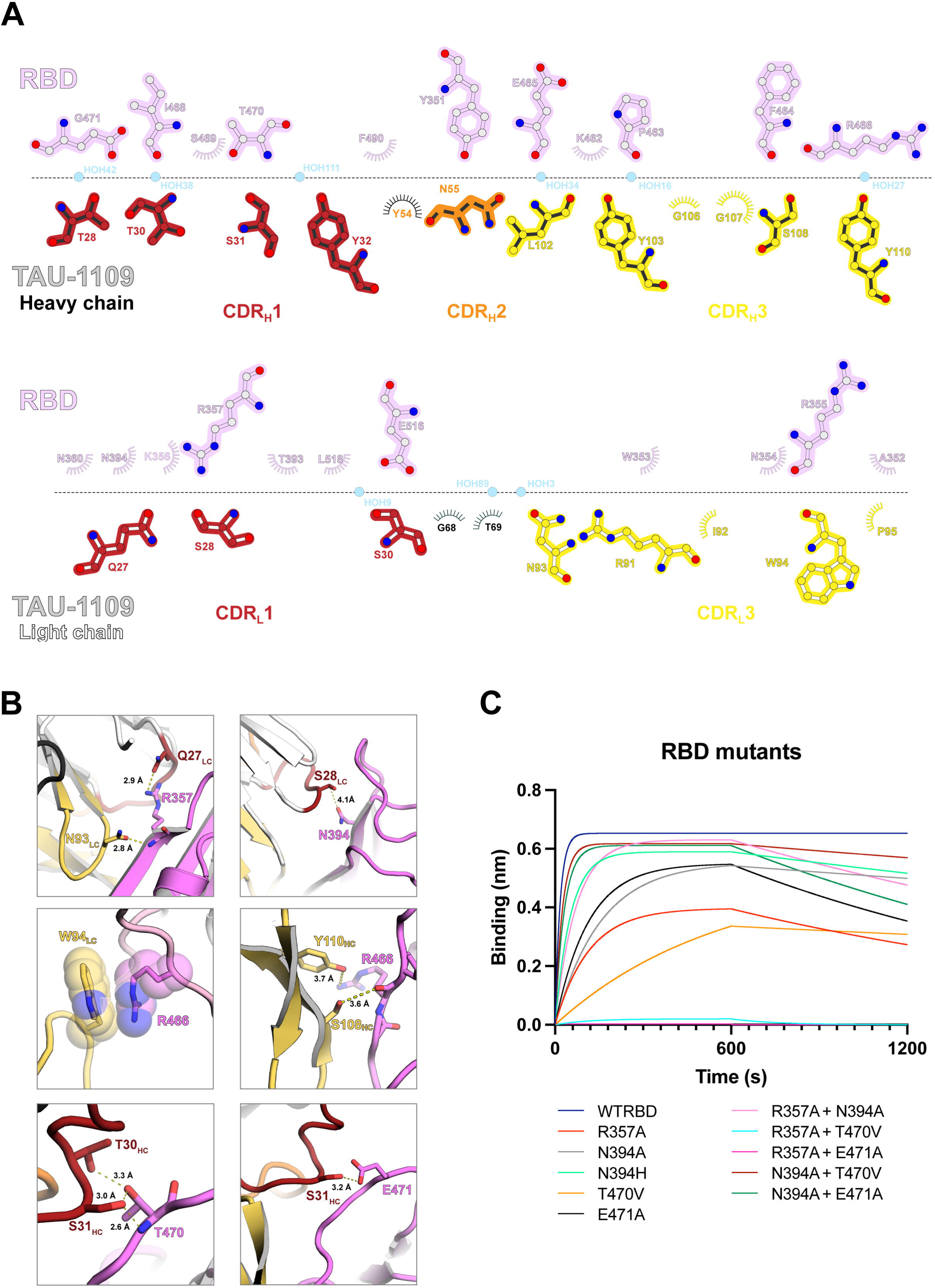
**The interactions between RBD and TAU-1109Fab** A) Ligplot representation ^47^ of the RBD-Fab^TAU-1109^ interface interactions. B) Detailed interactions between SARS-CoV-2 RBD residues (in violet) and respective CDRs of TAU-1109 Fab. The electron density (2Fo-Fc) map of all residues is displayed at a contour level of 1α over the mean value. The yellow dashes represent H-bonds between RBD and TAU-1109 Fab. C) Binding kinetics analysis of TAU-1109 with RBD mutants. WTRBD was used as a control. Ab TAU-1109 (10 μg/ml) was immobilized onto an anti-human Fc capture biosensor, followed by association with 200 nM RBD. The mutants are displayed in different colors.

**Table 2.**
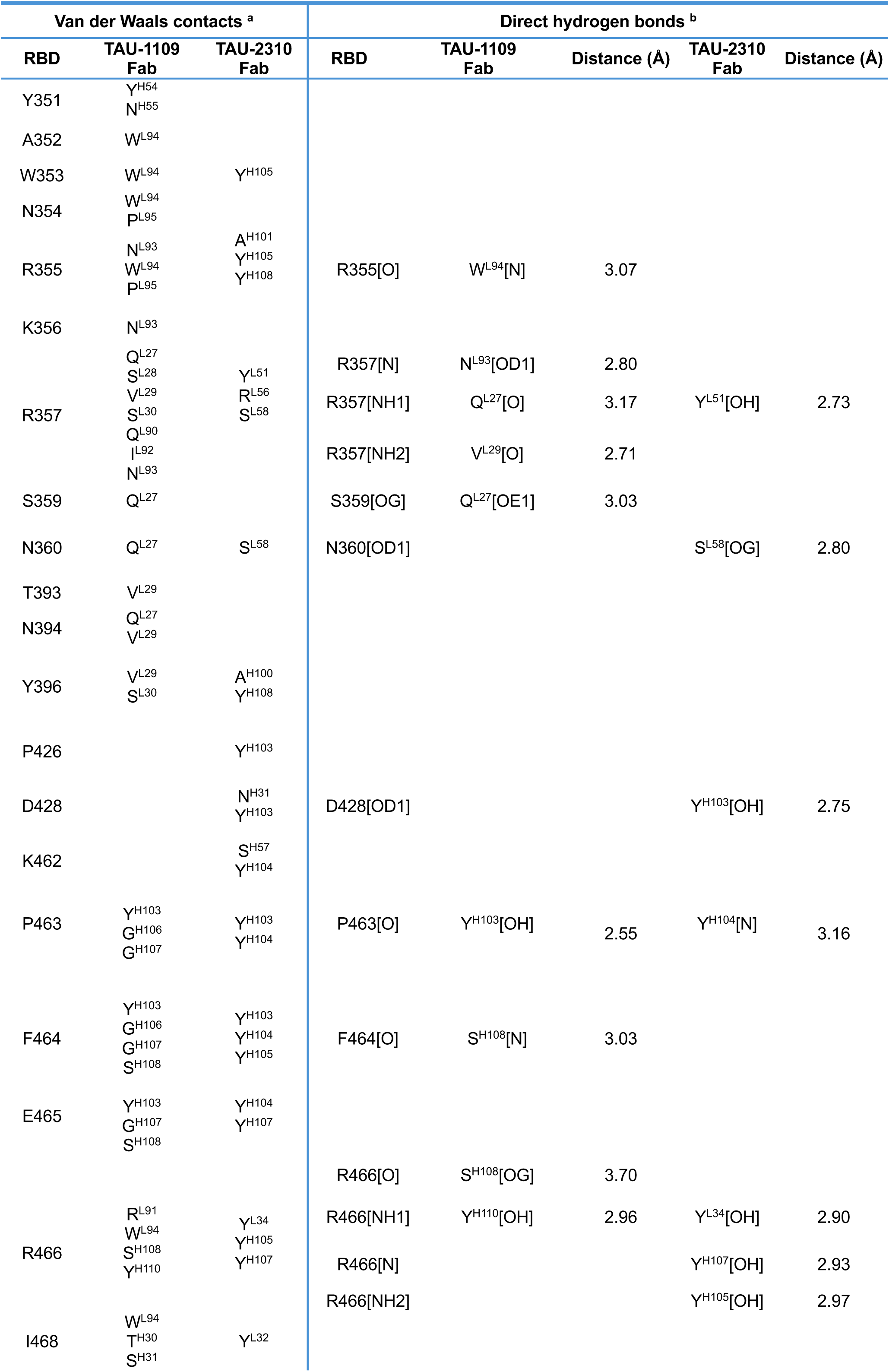
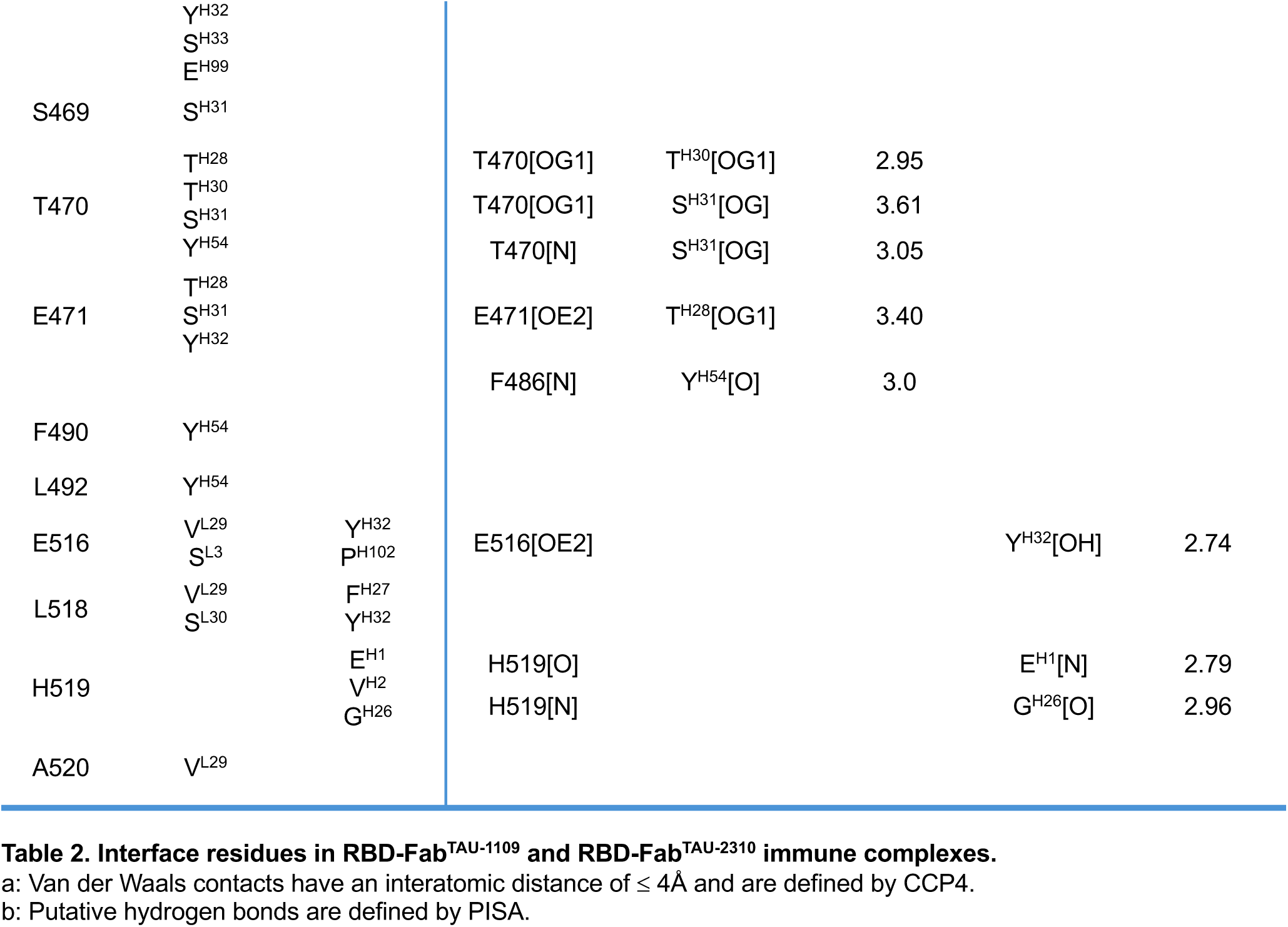
Interface residues in RBD-Fab^TAU-1109^ and RBD-Fab^TAU-2310^ immune complexes. a: Van der Waals contacts have an interatomic distance of ≤ 4Å and are defined by CCP4. b: Putative hydrogen bonds are defined by PISA.

The RBD-Fab^TAU-2310^ complex was crystallized in 0.2 M lithium citrate tribasic tetrahydrate and 20% (w/v) PEG 3350. The crystals appeared two days after drop setup and belonged to the *C2* space group. The structure was determined and refined at 2.12 Å (Table 1). The asymmetric unit of the RBD-Fab^TAU-2310^ complex crystals also contains one immune complex (Fig. 2A). The RBD-Fab^TAU-2310^ interface has a buried surface area of about 1211 Å^2^ and is composed of the residues W353, R355, R357, N360, Y396, P426, D428, K462, P463, F464, E465, R466, I468, E516, L518, and H519 in the RBD (Fig. 2C). Unlike RBD-Fab^TAU-1109^, the binding interface is smaller, and most interactions in the RBD-Fab^TAU-2310^ interface are contributed by the TAU- 2310 Fab heavy chain. The interface is stabilized by 10 direct hydrogen bonds, contributed by R357, N360, D428, P463, R466, E516, and H519, and several Van- der-Waals contacts (Fig. 4A-B, Table 2).

**Fig 4.**
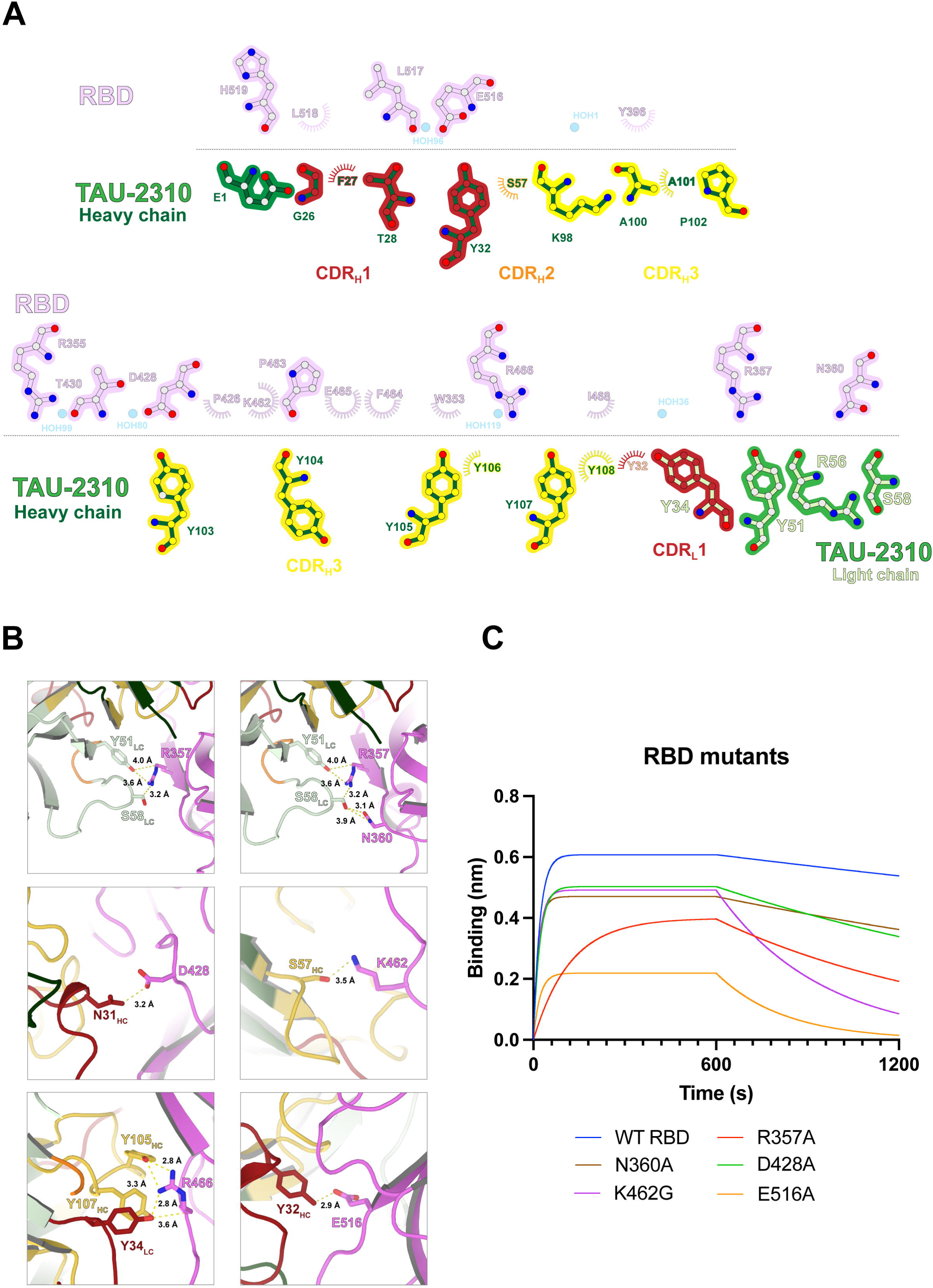
**The interactions between RBD and TAU-2310Fab** A) Ligplot representation ^47^ of the RBD-Fab^TAU-2310^ interface interactions. B) Detailed interactions between SARS-CoV-2 RBD residues (in violet) and respective CDRs of TAU-2310 Fab. The electron density (2Fo-Fc) map of all residues is displayed at a contour level of 1α over the mean value. The yellow dashes represent H-bonds between RBD and TAU-2310 Fab. C) Binding kinetics analysis of TAU-2310 with RBD mutants. WTRBD was used as a control. Ab TAU-2310 (10 μg/ml) was immobilized onto an anti-human Fc capture biosensor, followed by association with 200 nM RBD. The mutants are shown in different colors.

The epitopes of TAU-1109 and TAU-2310 do not overlap with the receptor-binding motif and are unlikely to sterically hinder the interaction between the RBD and ACE2, consistent with their lack of competition for hACE2 binding ^25^ (Fig. S1A). Most of the residues within the two epitopes are highly conserved across different VOCs of SARS-CoV-2 (Fig. S1B) and other viruses of the sarbecovirus subgenus (Fig. S1C), which likely explains the broad neutralization of the two antibodies across variants. To validate the interaction interfaces of each immune complex, we designed a panel of single and double RBD mutants and measured their interaction kinetics with the respective antibodies. The R357A, T470V, and E471A mutations caused a significant decrease in the binding affinities of TAU-1109 as compared to wild-type RBD (Fig. 3C, Table S2). In addition, the double mutants R357A-T470V and R357A-E471A demonstrated severe binding impairment (KD = 5.78 × 10⁻⁶ M and 5.28 × 10⁻⁷ M, respectively), indicating the critical role of these residues in the RBD-Fab^TAU-1109^ immune complex formation (Fig.3C, Table S2). Similarly, mutations within the TAU- 2310 epitope on the RBD (e.g., R357A, N360A, K462G, and E516A) resulted in a marked reduction in TAU-2310’s binding affinity (Fig.4C, Table S3). The conserved RBD residue, R466, interacts with W^L94^, S^H108^, Y^H110^ of TAU-1109 Fab and Y^L34^, Y^H105^, Y^H107^ of TAU-2310 Fab (Fig. 3B, 4B). Surprisingly, expression of RBD containing R466 mutations (e.g., R466A, R466E, R466D, R466C+A352C, and R466C+W353C) did not result in protein expression, suggesting that the residue R466 is crucial for RBD structural integrity.

### TAU-1109 and TAU-2310 target cryptic epitopes on the RBD and compete for binding in solution

Neutralizing antibodies against SARS-CoV-2 target different key regions on the spike protein, including the N-terminal domain (NTD), the stem helix, and the fusion peptide in the S2 subunit ^14,15,31,32^. However, most anti-SARS-CoV-2 antibodies discovered thus far target the receptor-binding domain (RBD) in the S1 subunit of the spike protein and have been classified into different classes (Fig. 5A) ^18,32^. While most classes of RBD-targeting antibodies are ineffective against the Omicron subvariants, the recently described classes 5 and 6 include antibodies that recognize cryptic, conserved regions in the RBD and exhibit broad neutralization capabilities ^23,33^. Notably, the binding of such antibodies has been demonstrated to induce greater conformational rearrangements in the spike trimers, contributing to their neutralizing abilities ^22,34,35^. While antibodies from classes 5 and 6 target cryptic epitopes in the RBD, there is no clearly defined demarcation between the boundaries of representative epitopes of the respective classes in the literature (Fig. 5A). The representative class 5 antibody, S2H97 (PDB ID: 7M7W), targets an epitope that is more closely positioned towards the binding epitope of class 4 nAbs, such as CR3022 (PDB ID: 6W41) (Fig. 5A) ^11,22^. Class 6 nAbs, on the other hand, are described to target the epitopes partway between class 3 and class 5 RBD-targeting antibodies (Fig. 5A) ^23^.

**Fig 5.**
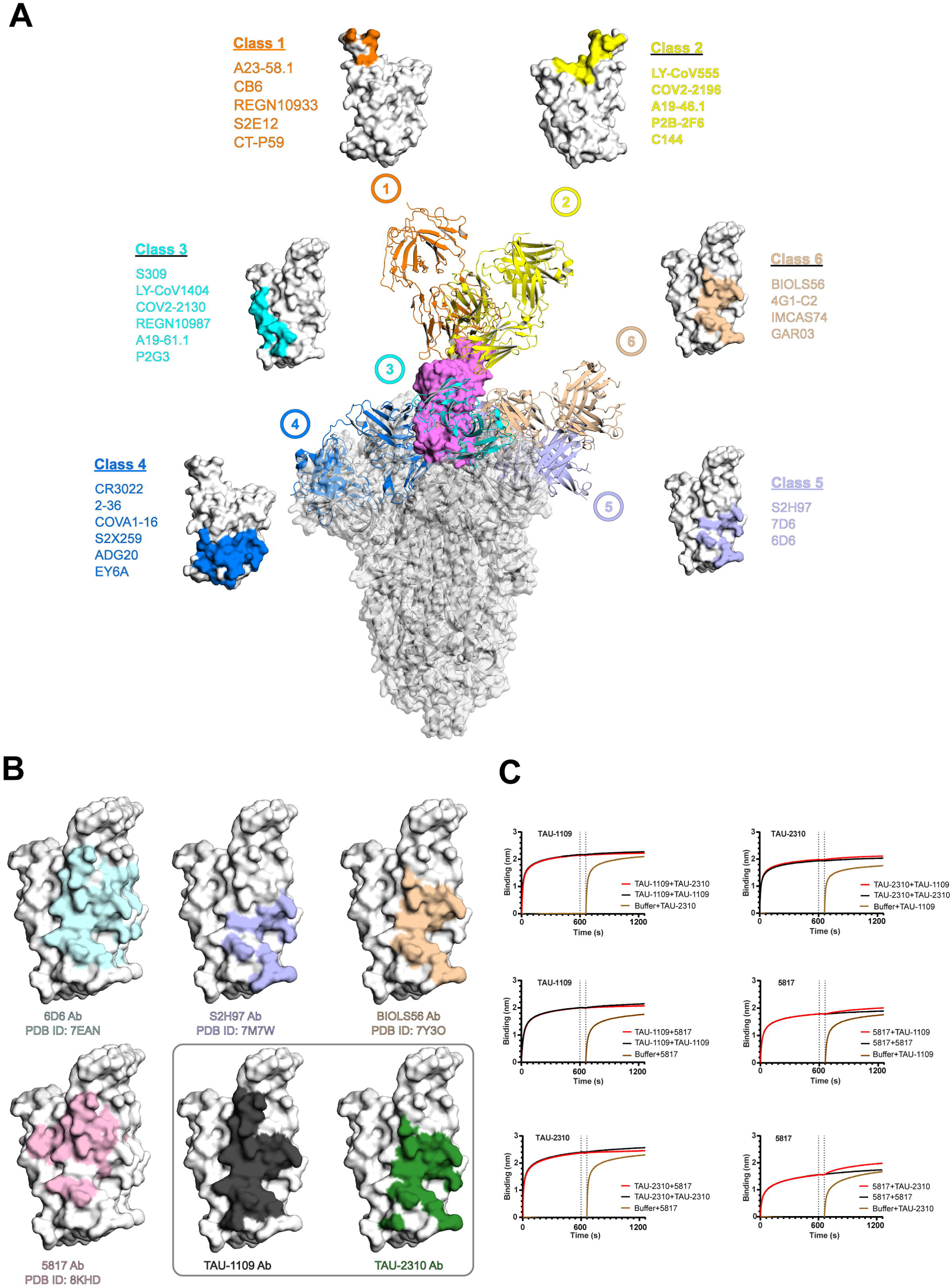
**While TAU-2310 targets an epitope similar to the class 6 nAbs, the TAU- 1109 epitope defines a unique structural footprint including residues from class 5 and class 6 RBD-neutralizing antibodies** A) Superimposition of structures of spike trimer (in grey) (PDB ID: 6VYB) and representative RBD-targeting neutralizing antibodies from each class (classes 1-6). The bound RBD is shown in violet. The epitopes of respective classes are shown separately on the RBD, and the nAbs belonging to the same class are listed in corresponding colors. B) Structural comparison of TAU-1109 and TAU-2310 epitopes with different class 5 and 6 neutralizing antibodies C) BLI-based binding competition assays. Both antibodies (200 nM) were sequentially associated for 600s each after a 60s baseline (shown by dotted lines) onto a biotinylated RBD loaded SAXS biosensor.

We compared the TAU-1109 and TAU-2310 epitopes with those targeted by different class 5 and 6 nAbs. Structural analysis revealed that TAU-2310 recognizes an epitope similar to that of other class 6 antibodies. However, TAU-1109 has a unique footprint that is widely comprised of residues common to the epitopes recognized by both classes 5 and 6 nAbs (Fig. 5B). Therefore, TAU-1109 can be classified as a class 6 antibody with an extended footprint (extended class 6 or class 6E), suggesting that both TAU-1109 and TAU-2310 may have the properties to conformationally destabilize the spike trimers upon binding, similar to other class 5 and 6 nAbs ^34,35^.

Considering the structural resemblance and partial overlap in the epitopes, we sought to determine whether TAU-1109 and TAU-2310 compete for the same binding site on the RBD in solution. To assess this, we performed a BLI-based competition assay, where each antibody was sequentially associated onto an RBD-loaded biosensor until saturation, followed by exposure to the second antibody. The results showed that TAU- 2310 was unable to bind to RBD-loaded biosensors after TAU-1109 association. In contrast, TAU-1109 showed slight binding to RBD pre-loaded with TAU-2310, indicating that TAU-1109 outcompetes TAU-2310 for binding to the RBD (Fig. 5C).

We further explored whether TAU-1109 and TAU-2310 compete with other known nAbs that target similar epitopes on the RBD. The recently reported broad neutralizing antibody mAb 5817, which targets a conserved epitope on the RBD, completely distinct from the ACE2 binding site, exhibits potent cross-variant neutralization of SARS-CoV-2 ^26^. Structural comparison of the mAb 5817, TAU-1109, and TAU-2310 epitopes revealed common residues participating in the formation of the respective immune complexes (Fig. 4B). We then performed the above-described competition assay to assess the binding efficacy of the 5817 mAb in the presence of TAU-1109 and TAU-2310, respectively. Our results show that mAb 5817 was unable to bind RBD pre-loaded with TAU-1109 or TAU-2310, whereas both TAU-1109 and TAU-2310 were able to efficiently bind to RBD pre-bound to mAb 5817 (Fig. 5C). Notably, the calculated buried surface area (BSA) in the RBD-5817 Fab immune complex is 979 Å^2 16^, which is significantly smaller than that of the RBD-Fab^TAU-1109^ (1902 Å^2^) and RBD- Fab^TAU-23^^10^ complexes (1211 Å^2^). Together, these results support our structural findings indicating that TAU-1109 Ab has a larger structural footprint and accordingly a higher binding affinity for the RBD as compared to other class 5 and 6 RBD-targeting antibodies, which may contribute to its elite neutralization potency and breadth (Fig. 5C).

### TAU-1109 and TAU-2310 binding induces a spatial clash to the SARS-CoV-2 spike trimer, resulting in antibody-mediated premature S1 shedding

The interaction of the SARS-CoV-2 RBD with the host ACE2 receptor destabilizes the S1–S2 interface, which promotes the S1 dissociation and induces irreversible conformational changes in the S2 subunit, ultimately driving its transition into the fusogenic state to mediate viral–host membrane fusion ^5^. While class 1–4 neutralizing antibodies (nAbs) inhibit viral entry by either directly obstructing the RBD- ACE2 interface or through steric hindrance ^10–12^, most class 5-6 RBD-targeting nAbs neutralize SARS-CoV-2 by triggering premature S1 shedding, rendering the spike protein non-functional for viral entry ^34^. To explore the effect of TAU-1109 and TAU- 2310 binding on the stability of spike trimer, we structurally modeled the RBD-Fab^TAU-1109^ and RBD-Fab^TAU-2310^ immune complexes onto the spike trimer, in both ‘open’ (PDB ID: 6VYB) and ‘closed’ (PDB ID: 6VXX) RBD conformations, which revealed a steric clash between the bound Fab ^TAU-1109/TAU-2310^ and the NTD of the adjacent protomer (Fig. 6). This finding suggests that the TAU-1109 and TAU-2310 epitopes, located distally from the RBM, reside on an RBD surface that faces the neighbouring NTD and is, otherwise, buried within the trimeric spike. This spatial constraint may underlie TAU- 1109 and TAU-2310’s ability to induce destabilization of the spike complex and subsequent premature S1 shedding.

**Fig 6.**
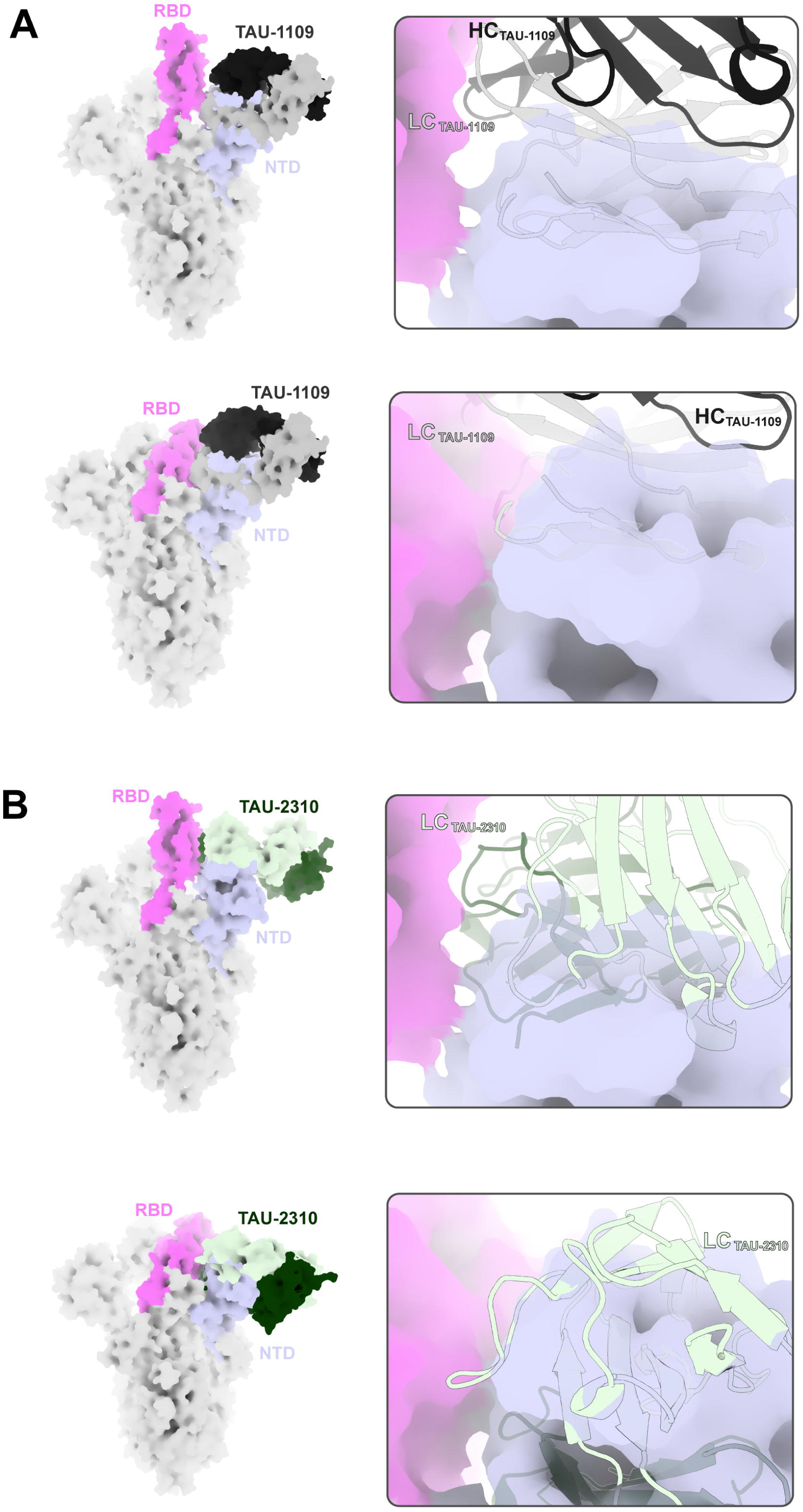
The TAU-1109 and TAU-2310 binding induces a spatial clash in the SARS- CoV-2 spike trimer. A) (Left) Superimposition of RBD-Fab^TAU-1109^ immune complex on spike trimer with RBD in ‘open’ (PDB ID: 6VYB) (top panel) or ‘closed’ (PDB ID: 6VXX) (bottom panel) conformations. On the right side, a zoom-in of the spatial clash between TAU-1109 Fab and the adjacent NTD protomer (light blue) of the spike trimer is shown for both ‘open’ (top panel) and ‘closed’ (bottom panel) RBD conformations. **B)** (Left) Superimposition of RBD-Fab^TAU-2310^ immune complex on spike trimer with RBD in ‘open’ (PDB ID: 6VYB) (top panel) or ‘closed’ (PDB ID: 6VXX) (bottom panel) conformations. On the right side, a zoom-in of the spatial clash between TAU-2310 Fab and the adjacent NTD protomer (light blue) of the spike trimer is shown for both ‘open’ (top panel) and ‘closed’ (bottom panel) RBD conformations.

To confirm our hypothesis, we designed an *in vitro* S1–S2 shedding assay to test the effect of TAU-1109 and TAU-2310 binding on the spike’s stability. HEK293T cells overexpressing the wild-type SARS-CoV-2 spike protein were incubated with TAU- 1109 and TAU-2310, respectively, and the extent of spike shedding was quantified over time using flow cytometry (Fig. S2). Interestingly, both TAU-1109 and TAU-2310 induced up to 50% spike shedding following 120 minutes of incubation, whereas CR3022, a class 4 antibody which is known to induce physical disruption of the spike trimer ^16^, did not trigger detectable S1 shedding effect (Fig. 7A). Furthermore, S1 shedding is expected to result in the release of soluble S1 subunit, presumably bound to the shedding antibody (TAU-1109/TAU-2310). To confirm this, we performed western blot analysis on cell lysates and culture supernatants from HEK293T cells overexpressing SARS-CoV-2 spike on their surface, following incubation with TAU- 1109 and or TAU-2310, respectively, at different time points. Cell lysates and supernatants were probed with an anti-S1-specific antibody, which revealed a distinct ∼ 50 kDa band in the supernatants corresponding only to the cells incubated with TAU-1109 or TAU-2310, confirming TAU-1109/TAU-2310-induced premature S1 shedding (Fig. 7B).

**Fig 7.**
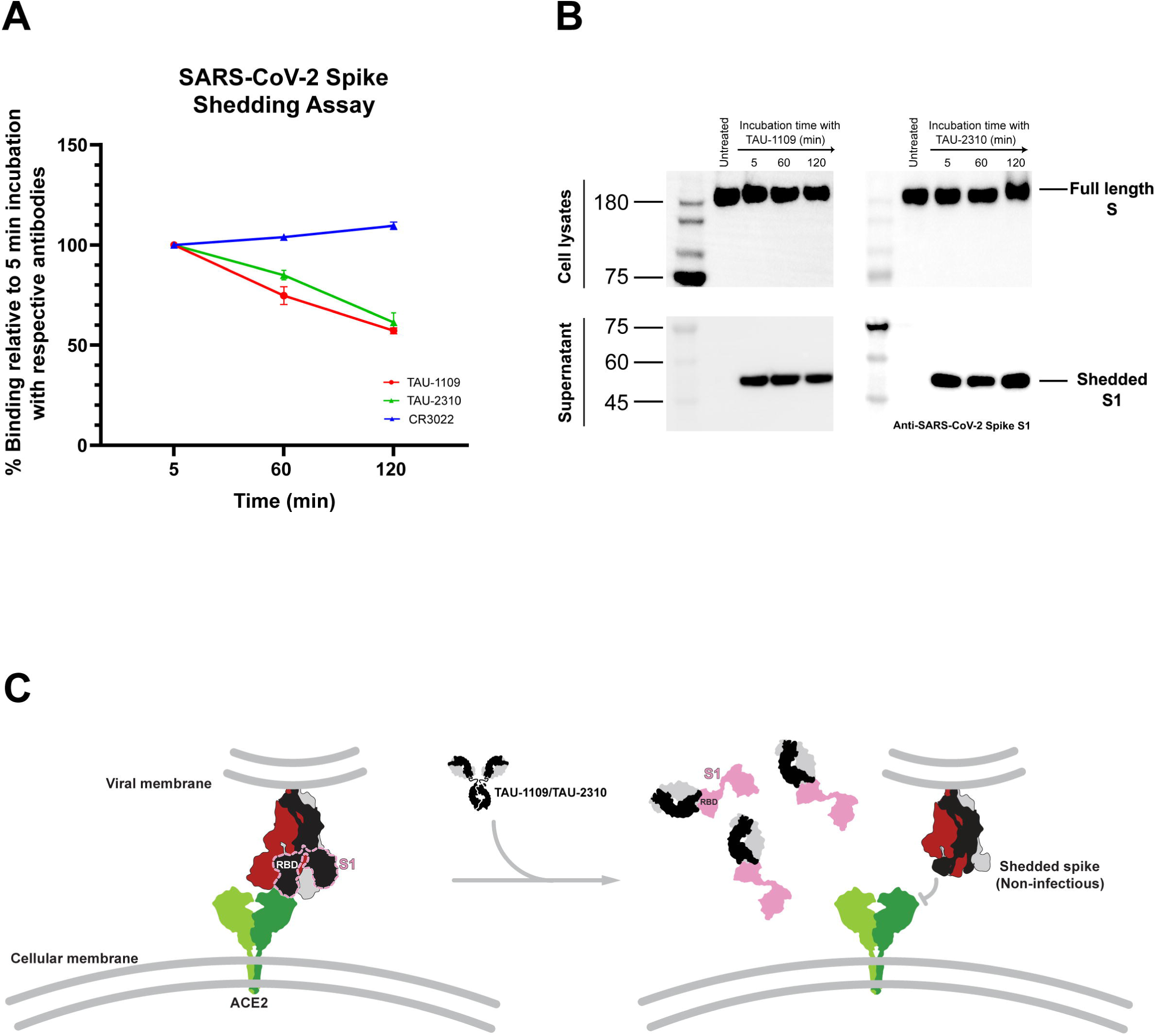
**The binding of Ab TAU-1109 and TAU-2310 to spike glycoprotein triggers premature S1 shedding** A) SARS-CoV-2 spike shedding assay based on flow cytometry. Ab CR3022 was used as a control. The data shown here were from three independent experiments, represented as mean±SEM. B) Western blot analysis of spike-transfected HEK 293T cell lysates and supernatants after serial incubation with TAU-1109 and TAU-2310. C) Possible mechanism of neutralization of TAU-1109 and TAU-2310. The TAU- 1109/TAU-2310 binding triggers premature S1 shedding, rendering the deformed spike incapable of host cell membrane fusion and subsequent infection.

Collectively, these findings suggest that both TAU-1109 and TAU-2310 bind to their cryptic epitopes on the RBD, stabilizing them in an exposed conformation that prevents their return to the native state. These interactions destabilize the spike trimer, promoting premature S1 shedding, and rendering the spike protein structurally compromised and incapable of mediating host cell entry and infection (Fig. 7C).

## Discussion

Since the onset of the COVID-19 pandemic, one of the most substantial challenges in the global response to SARS-CoV-2 infection has been the unprecedented rapid evolution of its genome, particularly within recent Omicron sub-lineages, which harbor novel spike protein mutations that confer extensive immune evasion capabilities ^21,29,36^. Most RBD-targeting neutralizing antibodies, isolated during the early stages of the pandemic, target mutation-sensitive epitopes in the RBM and have been proven ineffective against these newer variants ^28,30^. Therefore, the development of potent therapeutic monoclonal antibodies with broad neutralizing breadth has become a pressing priority to mitigate the severe consequences of SARS-CoV-2 infection.

Numerous neutralizing antibodies have been reported so far against the SARS-CoV- 2 spike protein ^18,32^, most of which target the RBD in the spike’s S1 subunit ^18,32^. These RBD-targeting neutralizing antibodies have been classified into different classes based on their targeted epitope and mode of binding to the spike protein ^37^.

*Class 1* RBD-targeting antibodies identify epitopes that overlap with the RBM, preventing the binding of the spike protein to the ACE2 receptor ^22,38–40^. Usually, these antibodies can recognize the RBD only in its ‘open’ (up) conformation. Since the RBM is highly prone to mutations during SARS-CoV-2 evolution, class 1 neutralizing antibodies have been found to lose their effectiveness against later-emerging VOCs compared to the original wild-type strain^19^. The *class 2* RBD-targeting antibodies also recognize the RBM, but unlike class 1 antibodies, they can bind to the RBD in both ‘open’ and ‘closed’ conformations^10,41,42^. Similar to class 1 antibodies, these class 2 antibodies also lack broad neutralization activity against many SARS-CoV-2 variants ^10^. In contrast, *class 3* RBD-targeting antibodies bind outside the RBM and can recognize the RBD regardless of its conformational state, often displaying broader neutralization across sarbecoviruses^12,43,44^. Similarly, *class 4* RBD-targeting antibodies recognize conserved RBD residues but do not directly block ACE2 binding ^11,45^. Instead, they interfere with ACE2 attachment through steric hindrance. Nonetheless, the mutations in newly emerged Omicron subvariants (like BQ.1.1 and XBB.1.5) lie in the binding epitopes of all four classes of nAbs, leading to a considerable escape of these variants from antibody neutralization ^21^. Recently, novel classes of RBD-targeting antibodies (designated classes 5 and 6) have been identified ^33–35^. These antibodies recognize highly conserved cryptic epitopes within the RBD that are typically obscured in the context of the intact spike trimer. Importantly, this binding specificity suggests their potential for broad neutralization across diverse SARS-CoV-2 variants and related sarbecoviruses ^33^.

The present study features two monoclonal antibodies (mAbs), TAU-1109 and TAU- 2310, isolated from two severe convalescent human patients eliciting a robust SARS-CoV-2-specific immune response. Structural and mechanistic investigations of TAU- 1109 and TAU-2310 revealed that these mAbs confer broad neutralization activity against different SARS-CoV-2 VOCs by targeting highly conserved cryptic epitopes on the receptor binding domain. Recent studies have reported different mAbs such as S2H97, 6D6, 7D6, 5817, and BIOLS56 recognizing conserved epitopes outside the ACE2 binding site ^22,26,34,35^. These antibodies effectively neutralize circulating VOCs by engaging unique epitopes beyond the mutation-prone RBM. Their ability to retain neutralizing activity across diverse variants reinforces the potential of conserved RBD epitopes as promising targets for next-generation therapeutic antibodies and universal pan-coronavirus vaccine development.

Despite targeting the non-ACE2 binding site in the spike protein, TAU-1109 and TAU- 2310 effectively neutralize SARS-CoV-2 infection *in vitro*. Previous studies have reported that some non-RBM targeting antibodies, such as CR3022 ^16^ and S2H97 ^22^, can induce premature S1 shedding. A similar phenomenon has also been documented in hepatitis E virus (HEV), where spatial clash was observed upon the binding of nAb 8C11 to virus-like particles, leading to the viral neutralization by antibody-induced physical disruption of the virions ^46^. In this study, the *in vitro* S1 shedding assay and western blot analysis confirmed the disruption of the S1 subunit after serial incubation of spike-transfected HEK 293T cells with TAU-1109/TAU-2310 mAbs. Hence, the structural and mechanistic insights gained from our study provide a comprehensive understanding of how naturally elicited antibodies engage the spike glycoprotein and destabilize its functional architecture. The identification of cryptic epitopes, which are not normally exposed but become accessible upon structural rearrangements following transient encounters with TAU-1109 and TAU-2310, offers a unique target space for the development of next-generation therapeutics and vaccines. Hence, this study highlights the potential of targeting conserved, cryptic epitopes on the SARS-CoV-2 spike protein to achieve broad, cross-variant neutralization. The broad-spectrum neutralization abilities of TAU-1109 and TAU-2310 emphasize the ongoing necessity for rigorous surveillance of SARS-CoV-2 variants to identify emerging mutations that might impact the prophylactic and therapeutic effectiveness of the monoclonal antibodies. It also underscores the importance of adopting a comprehensive approach to achieve broad-spectrum immunity by combining a diverse range of neutralizing antibodies that target various epitopes on the spike protein, as we showed in our previous study for TAU-1109 ^24^. Hence, in combination with our current findings, it is suggested that testing the efficacy of TAU-1109 or TAU-2310 in clinical settings can provide valuable insights into their potential as a targeted immunological therapy against SARS-CoV-2.

## Data Availability

Atomic coordinates and structure factors for the reported crystal structures have been deposited with the Protein Data bank under accession numbers 9SAT and 9SBB (RBD-Fab^TAU-1109^ and RBD-Fab^TAU-2310^ immune complexes, respectively).

## Author contributions

A.H. and M.D. planned and performed the biochemical experiments, analysed data, prepared the figures, and wrote the manuscript, M.M and N.T.F. produced RBD mutants and RBD VOC, R.Y. and N.T.F. produced the 5817 nAb, L.S.I.T. subcloned and produced stable cell lines expressing wtRBD, M.D. and M.G.T. established the pseudo-viral assays, M.D. and N.T.F conceptualized the study. All authors proofread the manuscript.

## Acknowledgements

We would like to thank the staff of beamlines ID30B and ID23-2 at the European Synchrotron Radiation Facility (ESRF, Grenoble, France) for their assistance in data collection and technical support of the beamline. This work was supported by the Israel Science Foundation (ISF) grants [401/18] and [352/23] to MD, grants [3136/22] and [638/23] to NTF; Binational Science Foundation (BSF) [01031771] to NTF; BMGF INV- 058519 to NTF. RY was supported by a PhD Scholarship from the Tel Aviv University Center for Combatting Pandemics.

