## Supplementary material for "Monoclonal Antibodies from COVID-19 Convalescent Patients Target Cryptic Epitopes for Universal SARS-CoV-2 Neutralization": Supplenental figures and tables

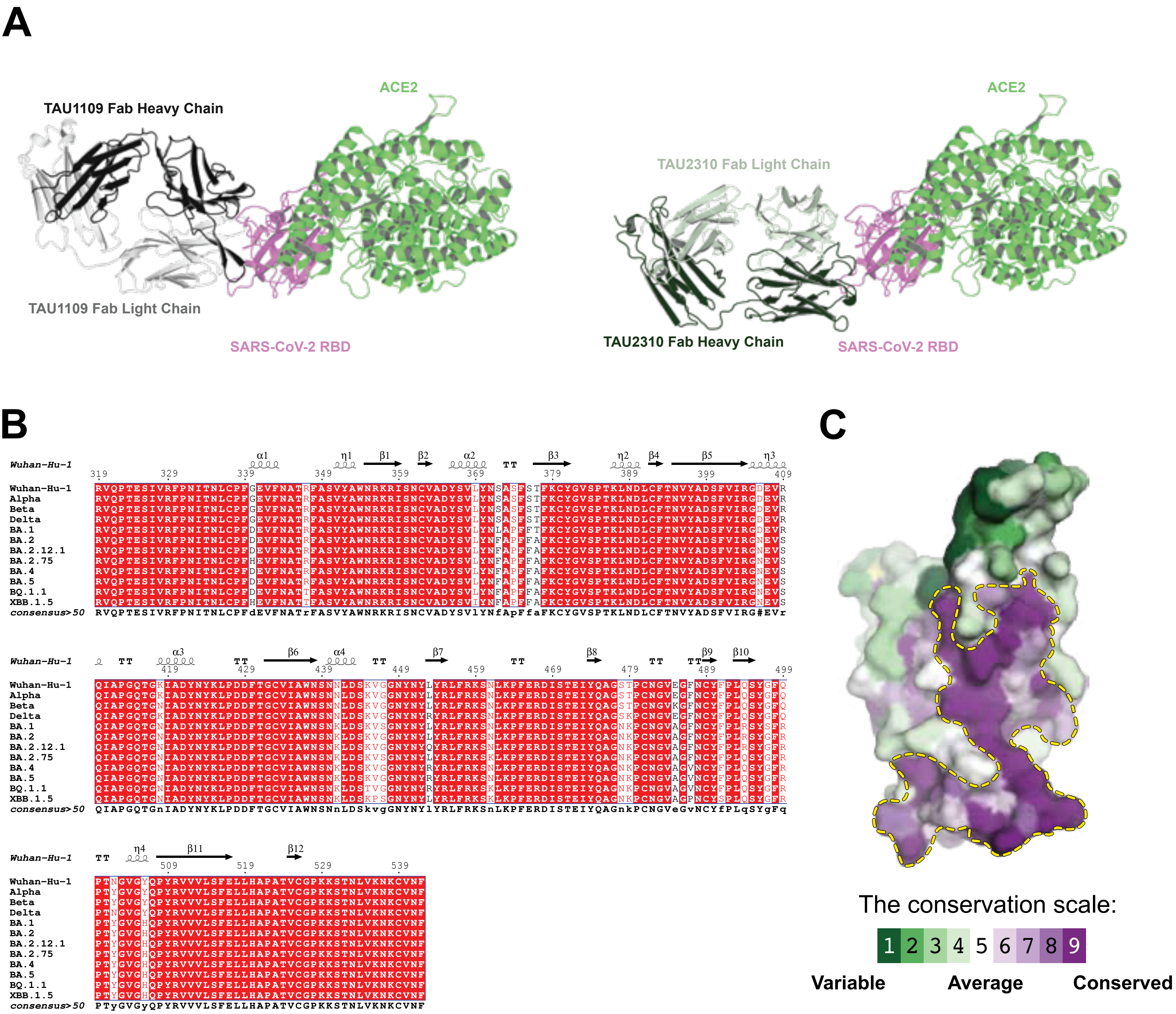


**Fig S1. Epitope orientation and RBD conservation analysis. (A)** The cartoon representation showing TAU1109 Fab (left) and TAU2310 Fab (right) and ACE2 binding (lime green) on SARS-CoV-2 RBD showing that TAU-1109 and TAU2310 do not interfere with ACE2 binding. **(B)** Sequence alignment ofRBDs of different VOCs of SARS-CoV-2. The secondary structure prediction is depicted according to RBD structure in wtRBD-FabTAU1109 immune complex crystal structure. The plot was prepared using an online server (https://espript.ibcp.fr/ESPript/ESPript/). **(C)** Surface represention of wtRBD colored according to conservation scores of residues across Sarbecovirus subgenus as determined by Consurf webserver.

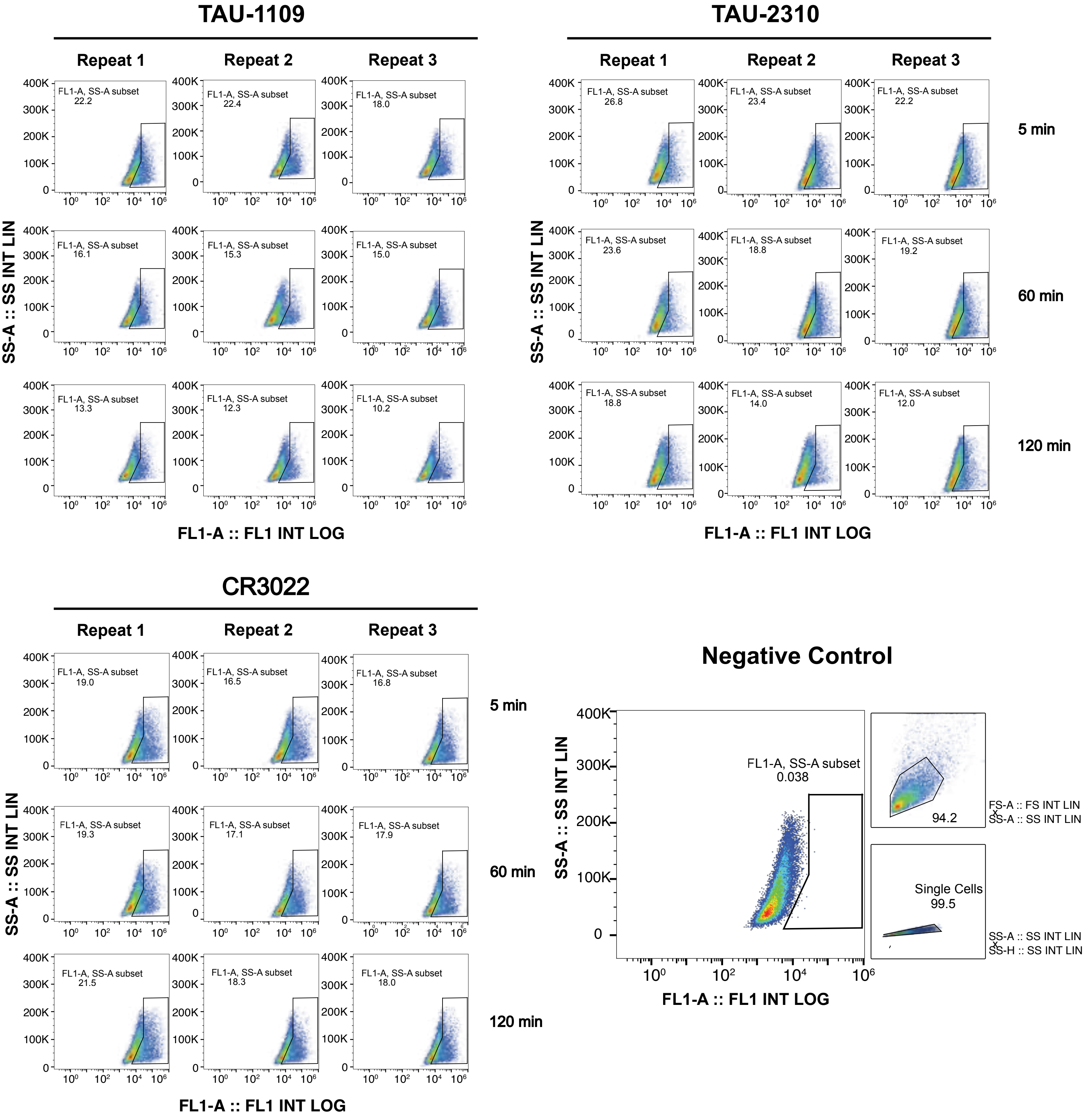


**Fig S2. The Ab-induced S1 shedding was measured by an in vitro S1/S2 shedding assay based on flow cytometry.** The HEK293T cells expressing the wild-type spike on the surface were incubated with the respective antibodies for different time points. The percentage of cells at each time point was determined by the number of positive cells in the selected gates. Ab CR3022 was used as a control.

### Table S1. Binding affinity kinetics of TAU-1109 and TAU-2310 against the RBDs of different VOCs of SARS-CoV-2

| | | *Wuhan-Hu-1* | *B.1.1.529* | *BA.2* | *BA.2.12.1* | *BA.2.75* | *BA.4.6* | *BQ.1.1* | *XBB.1.5* |
|---|---|---|---|---|---|---|---|---|---|
| ***TAU-1109*** | $k_a$ (1/Ms) | 2.01E+05 | 1.05E+05 | 1.52E+05 | 1.53E+05 | 2.44E+05 | 2.23E+05 | 3.20E+05 | 1.81E+05 |
| | $k_{dis}$ (1/s) | 1.35E-04 | 9.14E-05 | 1.48E-04 | 7.30E-05 | 9.56E-05 | 4.15E-04 | 4.96E-04 | 8.30E-04 |
| | **$K_D$ (M)** | **6.73E-10** | **8.72E-10** | **9.76E-10** | **4.78E-10** | **3.92E-10** | **1.86E-09** | **1.55E-09** | **4.58E-09** |
| ***TAU-2310*** | $k_a$ (1/Ms) | 2.34E+05 | 3.90E+05 | 2.10E+05 | 3.26E+05 | 1.76E+05 | 1.62E+05 | 1.92E+05 | 1.69E+05 |
| | $k_{dis}$ (1/s) | 2.03E-04 | 2.53E-04 | 1.82E-04 | 1.75E-04 | 8.01E-05 | 2.71E-04 | 3.11E-04 | 2.31E-04 |
| | **$K_D$ (M)** | **8.67E-10** | **6.49E-10** | **8.69E-10** | **5.38E-10** | **4.55E-10** | **1.67E-09** | **1.62E-09** | **1.37E-09** |

### Table S2. Binding affinity kinetics of TAU-1109 against various RBD mutants

| | | **R357A** | **N394A** | **N394H** | **T470V** | **E471A** | **R357A + N394A** | **R357A + T470V** | **R357A + E471A** | **N394A + T470V** | **N394A + E471A** |
|---|---|---|---|---|---|---|---|---|---|---|---|
| **TAU-1109** | $k_a$ (1/Ms) | 4.08E+04 | 3.18E+04 | 9.35E+04 | 8.29E+03 | 4.11E+04 | 6.85E+04 | 1.22E+04 | 4.64E+04 | 1.48E+05 | 8.14E+04 |
| | $k_{dis}$ (1/s) | 6.16E-04 | 1.39E-04 | 2.22E-04 | 1.45E-04 | 7.28E-04 | 4.68E-04 | 6.44E-03 | 2.68E-01 | 8.87E-05 | 5.73E-04 |
| | **$K_D$ (M)** | **1.51E-08** | **4.38E-09** | **2.37E-09** | **1.74E-08** | **1.77E-08** | **6.84E-09** | **5.28E-07** | **5.78E-06** | **6.01E-10** | **7.04E-09** |

### Table S3. Binding affinity kinetics of TAU-2310 against various RBD mutants

| | | **R357A** | **N360A** | **D428A** | **K462G** | **E516A** |
|---|---|---|---|---|---|---|
| **TAU-2310** | $k_a$ (1/Ms) | 3.58E+04 | 3.07E+05 | 2.30E+05 | 2.60E+05 | 2.14E+05 |
| | $k_{dis}$ (1/s) | 1.21E-03 | 4.35E-04 | 6.59E-04 | 2.91E-03 | 4.49E-03 |
| | **$K_D$ (M)** | **3.37E-08** | **1.42E-09** | **2.87E-09** | **1.12E-08** | **2.10E-08** |

**Table S4. Mutagenesis primers for various RBD mutants based on the TAU-1109 and TAU-2310 epitope**

| Primers | Sequence (5' to 3') |
|---|---|
| RBD_A352C_F | TGCCTCTGTCTATTGCTGGAACAGGAAGAG |
| RBD_A352C_R | CTCTTCCTGTTCCAGCAATAGACAGAGGCA |
| RBD_D428A_F | AACTGCCTGATGCGTTCACAGGCTGTGT |
| RBD_D428A_R | ACACAGCCTGTGAACGCATCAGGCAGTT |
| RBD_E516A_F | GCTGTCCTTTGCGCTGCTCCATG |
| RBD_E516A_R | CATGGAGCAGCGCAAAGGACAGC |
| RBD_K462G_F | AAGAGCAACCTGGGCCCATTTGAGAG |
| RBD_K462G_R | CTCTCAAATGGGCCCAGGTTGCTCTT |
| RBD_N360A_F | AAGAGGATTAGCGCGTGTGTGGCTGACTAC |
| RBD_N360A_R | GTAGTCAGCCACACACGCGCTAATCCTCTT |
| RBD_R357A_F | CTGGAACAGGAAGGCGATTAGCAACTGTGT |
| RBD_R357A_R | ACACAGTTGCTAATCGCCTTCCTGTTCCAG |
| RBD_R466C_F | ACCATTTGAGTGCGACATCAGCACAGA |
| RBD_R466C_R | TCTGTGCTGATGTCGCACTCAAATGGT |
| RBD_W353C_F | CTGTCTATGCCTGCAACAGGAAGAGGAT |
| RBD_W353C_R | ATCCTCTTCCTGTTGCAGGCATAGACAG |
| RBD_394A_F | CTCAATGATTTGTGTTTCACGgcTGTATACGCCGATTCCTTC |
| RBD_394A_R | GAAGGAATCGGCGTATACAgcCGTGAAACACAAATCATTGAG |
| RBD_394H_F | CTCAATGATTTGTGTTTCACGcATGTATACGCCGATTCCTTC |
| RBD_394H_R | GAAGGAATCGGCGTATACATgCGTGAAACACAAATCATTGAG |
| RBD_T470V_F | GAACGCGACATTTCTgtCGAAATATATCAAGCCGGC |
| RBD_T470V_R | GCCGGCTTGATATATTTCGacAGAAATGTCGCGTTC |
| RBD_E471A_F | CGCGACATTTCTACCGcAATATATCAAGCCGGCAGTACG |
| RBD_E471A_R | CGTACTGCCGGCTTGATATATTgCGGTAGAAATGTCGCG |
| RBD_R466A_F | AACCTCAAGCCTTTTGAAgcCGACATTTCTACCGAAATATATC |
| RBD_R466A_R | GATATATTTCGGTAGAAATGTCGgcTTCAAAAGGCTTGAGGTT |
| RBD_R466D_F | AACCTCAAGCCTTTTGAAgaCGACATTTCTACCGAAATATATC |
| RBD_R466D_R | GATATATTTCGGTAGAAATGTCGtcTTCAAAAGGCTTGAGGTT |
| RBD_R466E_F | AACCTCAAGCCTTTTGAAgagGACATTTCTACCGAAATATATC |
| RBD_R466E_R | GATATATTTCGGTAGAAATGTCctcTTCAAAAGGCTTGAGGTT |
